# Mouse tissue harvest-induced hypoxia rapidly alters the in vivo metabolome, between-genotype metabolite level differences, and ^13^C-tracing enrichments

**DOI:** 10.1101/2022.06.07.495179

**Authors:** Adam J. Rauckhorst, Nicholas Borcherding, Daniel J. Pape, Alora S. Kraus, Diego A. Scerbo, Eric B. Taylor

## Abstract

Metabolism is potently regulated by oxygen as the terminal acceptor of the electron transport chain. Thus, a challenge for capturing the in vivo metabolome of animal tissues is to achieve rapid freezing after dissection-induced loss of perfusion before the onset of hypoxia-driven metabolomic remodeling. However, the timing of the metabolomic changes elicited by post-dissection freezing delays are not well described. We addressed this problem by carefully and systematically assessing broad, genotype-specific, and ^13^C isotopologue metabolomic change resulting from post-dissection, ex vivo mouse tissue metabolism. Based on experiments with mouse liver, heart muscle, and skeletal muscle, we show that broad metabolomic change is rapid, that both false negative and false positive between genotype differences are induced, and that ^13^C-isotopologue abundances and enrichment percentages change with post-dissection hypoxia. These findings provide a previously absent, systematic illustration of the extensive and confounding metabolomic changes occurring within the early minutes of delayed tissue freezing.

## Introduction

Metabolomics is a powerful investigative approach to solve biological problems that is dramatically increasing in use. The metabolome is the endogenous small molecule pool of reaction substrates and products (1). It links cellular chemistry, energetics, and environment to impact phenotype, responding nearly instantaneously to homeostatic challenges (2). Thus, the metabolome is a functional, real-time property of health and disease states. Metabolomics refers to the measurement of the metabolome using NMR or mass spectrometry, with the latter capable of measuring hundreds of metabolites in milligram quantities of sample (3). As evidence of metabolomics’ growing use, a PubMed.gov search for ‘metabolomics or metabolome’ at the time of this writing returned >55,000 original research articles published since the first search-retrievable, mention of “metabolome" in 1998 (4). Notably, greater than 50% of these manuscripts were published after 2017, with over 90% after 2011.

Because broad metabolomic analysis of animal tissues requires dissection, a major risk to the integrity of metabolomic data is a hypoxia-induced shift of metabolism, between the time of disrupted perfusion and freezing. Indeed, beginning with the evolution of complex life, cellular metabolism became programmed to be potently regulated by oxygen (5–8). In most eukaryotic cells, mitochondrial electron transport chain flux coupled to oxygen reduction regulates an array of redox-sensitive metabolic reactions and drives bulk ATP production. Furthermore, in addition to supplying nutrients and removing cellular waste, perfusion evolved to provide tissues with oxygen. Consequently, animal tissue metabolism is intricately connected to oxygen availability, and disrupted perfusion rapidly leads to evolutionarily imbedded, coordinated metabolic responses. Thus, to rigorously investigate the in vivo metabolome of animal tissues, it is essential to quickly freeze tissues after dissection to prevent ex vivo, hypoxia-driven metabolomic remodeling.

However, how delayed tissue freezing affects the broad metabolome during the immediate post-dissection period has remained unaddressed. Although it is well known that disrupted oxygen supply rapidly alters cellular energy charge, the speed and extent to which this extends to the larger metabolome has not been defined. Furthermore, most manuscripts reporting tissue metabolomic analysis do not precisely specify the lag time between loss of perfusion and tissue freezing. The absence of time to freeze information makes it difficult to infer how post-dissection metabolism may have impacted the reported metabolomic data. To address this problem, we performed experiments evaluating how the metabolome of three mouse tissues changes with delayed freezing after harvest. We show that tissue harvest-induced hypoxia rapidly alters the in vivo metabolome, between-genotype metabolite level differences, and ^13^C-tracing enrichments. With metabolomics being increasingly adopted as a mainstream investigative approach and the bloom of multiomics, this is an especially critical time for the field of metabolism to employ best practices to obtain reliable metabolomic data. Our findings provide a previously absent, systematic illustration of the extensive and confounding metabolomic changes occurring within the early minutes of delayed tissue freezing.

## Results

### Delayed post-dissection freezing of the liver leads to broad metabolomic drift

The liver is a metabolically versatile tissue routinely used to investigate basic and translational metabolism. To understand how quickly after dissection from live, anesthetized mice liver tissue must be frozen to preserve the in vivo metabolome, we performed time-course experiments followed by broad metabolomic analysis. Specifically, the left lateral lobe of the liver was dissected from isoflurane-anesthetized mice and frozen by liquid nitrogen-temperature freeze clamping at various timepoints after dissection: immediately (about 1 second), and after 30 seconds, 1 minute, 3 minutes, and 10 minutes (Figure 1A). Compared to immediate freezing, ex vivo metabolomic drift was rapid and progressive (Figure 1B, Table S1). Thirty-one metabolites significantly increased by 30 seconds after dissection, and 128 did so by 10 minutes (Figure 1C).

**Figure 1:**
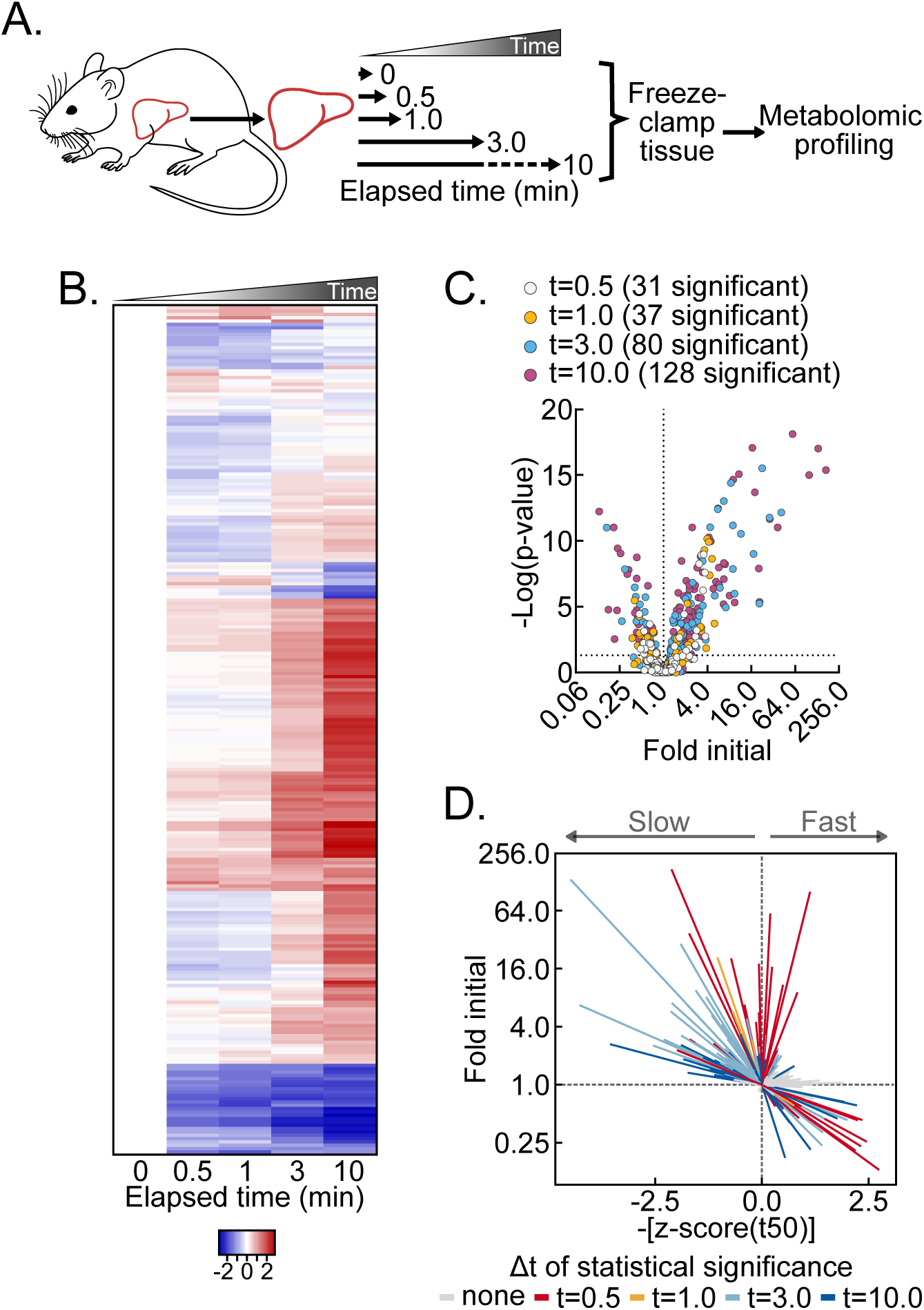
Post-dissection hypoxia rapidly alters the liver metabolome. (A) Schematic of experimental procedures. (B) Heatmap depicting the relative change in the metabolome observed following delayed freezing over the time course (all samples compared to t=0 samples). Data are presented as the mean/time point (n=5-6 biological replicates). (C) Volcano plot of metabolomic data shown in panel B. Log_2_ fold calculated for each time point relative to t=0. P-values were calculated one-way ANOVA with post-hoc Holm-Sidak multiple comparison test versus t=0 on log transformed data. The number of significant differences for each time point are listed in panel. (n=5-6 biological replicates). (D) Urchin plot where each line represents a single metabolite. X-axis shows the scaled time required to reach 50% metabolite change (t50) and the y-axis shows the Log_2_(Fold) abundance at 10 minutes. Line coloring denotes the time point at which the metabolite change becomes significant as calculated in panel (C).

Given these extensive changes across individual metabolites, we aimed to define broad properties of metabolomic change elicited by delayed freezing after liver tissue harvest. We calculated and averaged the time to 50% change for all metabolites. This produced a time of one-half fold-change (t50) for the broad metabolome of approximately 3.58 +/- 0.32 minutes (Figure S1A). We then plotted all measured metabolites as individual line vectors, by magnitude of change versus negative z-scores of t50s, with time-points of statistically significant change indicated by line color (Figure 1D, Table S2). This vectorized “urchin plot” displays line slopes as an interaction between rate of change kinetics and magnitude of change (Figure S1B). An interactive, fully labeled web object version of the urchin plot is available online (Object S1). Here, it reveals several features of dissection-induced metabolomic drift. Significantly changed metabolites binned to the upper left, upper right, lower right, but not lower left quadrants. This pattern indicates that, on average, metabolite decreases reach time of 50% change more rapidly than metabolite increases. The absence of metabolites in the lower left quadrant is consistent with larger macromolecules not detected by small molecule metabolomic analysis, like glycogen, being slowly depleted while feeding metabolic pathways that show hypoxic metabolite accumulation. Overall, these data show post-tissue dissection metabolomic drift is diverse in kinetics and magnitudes, with both observed metabolomic and likely non-observed macromolecular metabolic interactions.

### The post-dissection liver metabolome is rapidly remodeled by hypoxia

To begin evaluating changes in individual metabolites, we asked whether metabolites known to be influenced by hypoxia changed as expected during delayed tissue freezing. The ratio of the critical redox cofactor couple NAD+ and NADH is highly sensitive to hypoxia. NADH accumulation results from hypoxia because the mitochondrial electron transport chain becomes fully reduced in the absence of a terminal electron acceptor (Figure 2A). NADH levels were strikingly increased by 30 seconds, reaching a near peak at 2-fold initial levels with minimal further increases by 10 minutes (Figure 2B). This was accompanied by a concomitant decrease in NAD+ levels and, thus, an increased NADH:NAD+ ratio (Figure 2B, S2A). Without electron transport chain activity, lost mitochondrial inner membrane proton motive force results in decreased mitochondrial ATP synthesis. ATP levels significantly decreased by 30 seconds of tissue dissection and continued decreasing throughout 10 minutes (Figure 2C). In parallel, ADP modestly increased after 3 minutes, while AMP levels doubled by 30 seconds and accumulated to greater than 500% of initial levels after 10 minutes. As an additional control, we tested whether phosphorylation of the master energy-sensing kinase AMPK tracked with the AMP:ATP ratio (9–12). The AMP:ATP ratio increased nearly 3-fold by 30 seconds after dissection, and it continued to increase through 10 minutes (Figure S2B). In parallel, AMPK alpha subunit threonine 172 phosphorylation, which is known to increase when the AMP:ATP ratio increases, rose linearly through 3 minutes, reaching a nearly 700% increase by 10 minutes (Figure 2D). Thus, ex vivo tissue hypoxia following dissection quickly leads to a reduced redox balance and a decreased energy state.

**Figure 2:**
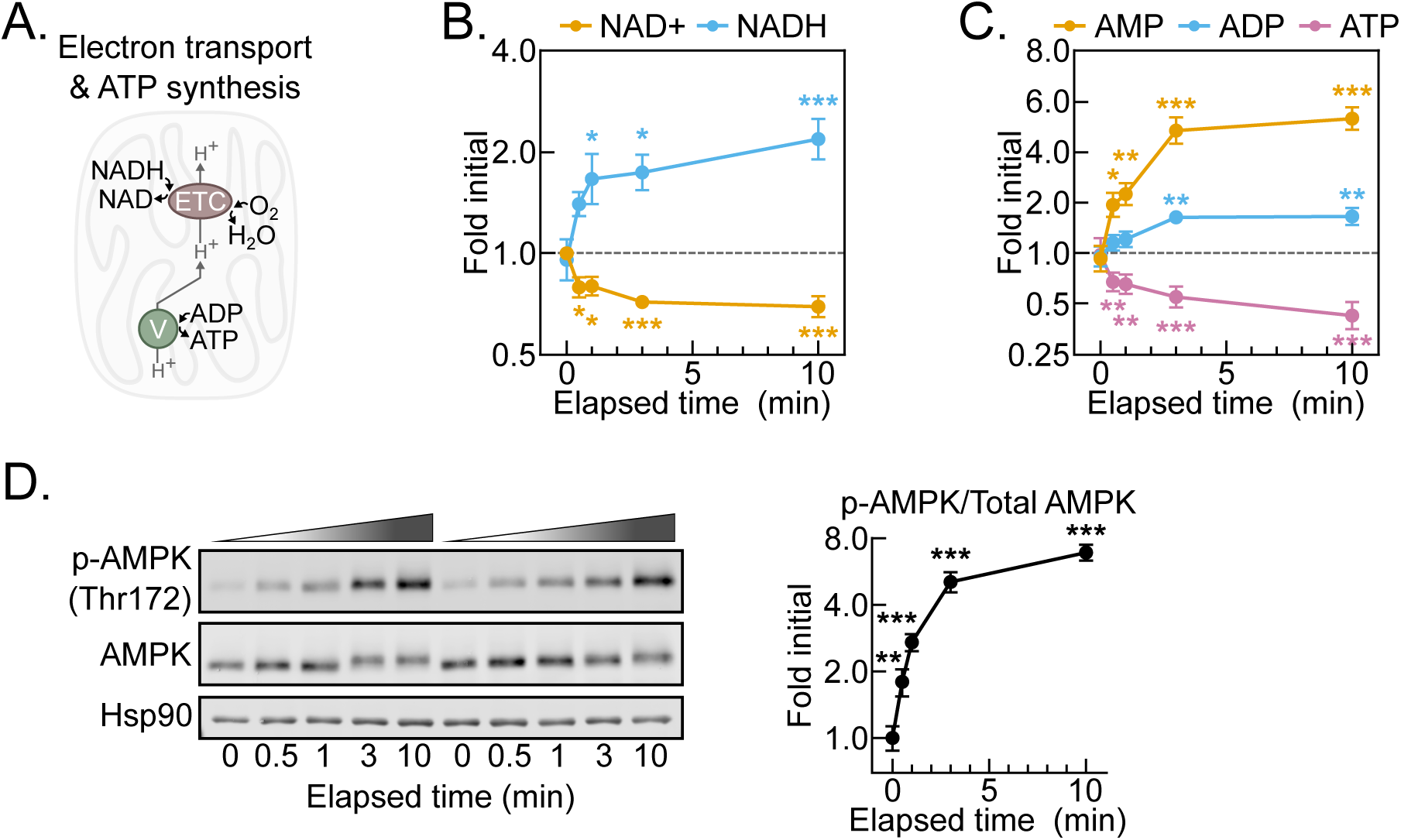
Post-dissection hypoxia results in NAD+ reduction and energetic crisis. (A) Schematic depicting dependence of NADH oxidation and O_2_ reduction activities of the electron transport chain (ETC) to the ATP synthesis activity of the ATP synthase/Complex V (V). (B-C) NAD+ and NADH (B) and AMP, ADP, and ATP (C) levels over time between dissection and freezing. Data are presented as the mean±SEM. Statistics were calculated using one-way ANOVA with post-hoc Holm-Sidak multiple comparison test versus t=0 on log transformed data. (n=6 biological replicates). (D) Representative western blot of p-AMPK (Thr172), AMPK, and Hsp90 of lysates generated from liver tissue collected throughout delayed freezing time course, and relative quantification of phosphorylated and total AMPK. Relative quantification data presented as the mean±SEM. Statistics calculated by one-way ANOVA with post-hoc Holm-Sidak multiple comparison test versus t=0 on log transformed data. (n=6 biological replicates). *p<0.05, **p<0.01, ***p<0.001; Related to Figure S1 and Tables S1 and S2.

### Post-dissection liver hypoxia leads to increased glycolytic and decreased TCA cycle activity

We next considered the rate at which a reduced redox balance and decreased energy state propagate to changes in central carbon metabolism metabolites. We examined metabolites in glycolysis and the TCA cycle, which are regulated by the NADH:NAD+ and AMP:ATP ratios (Figure 3A). Indeed, a hypoxic shift to increased glycolytic and decreased TCA cycle activity throughout the time-course is evident from a final, greater than 20-fold increase in the lactate:citrate ratio (Figure S3A). In accord, most glycolytic intermediates increased, except for 3-phosphoglycerate and pyruvate, likely because of relatively increased consumption to support ATP production and NAD+ regeneration, respectively (Figures 3A, 3B). Conversely, but also consistent with an elevated NADH:NAD+ ratio, TCA cycle intermediates decreased, except for succinate. Notably, succinate is a well-recognized marker of hypoxia and increased 9-fold by 10 minutes, which further validates our approach (Figure 3C).

**Figure 3:**
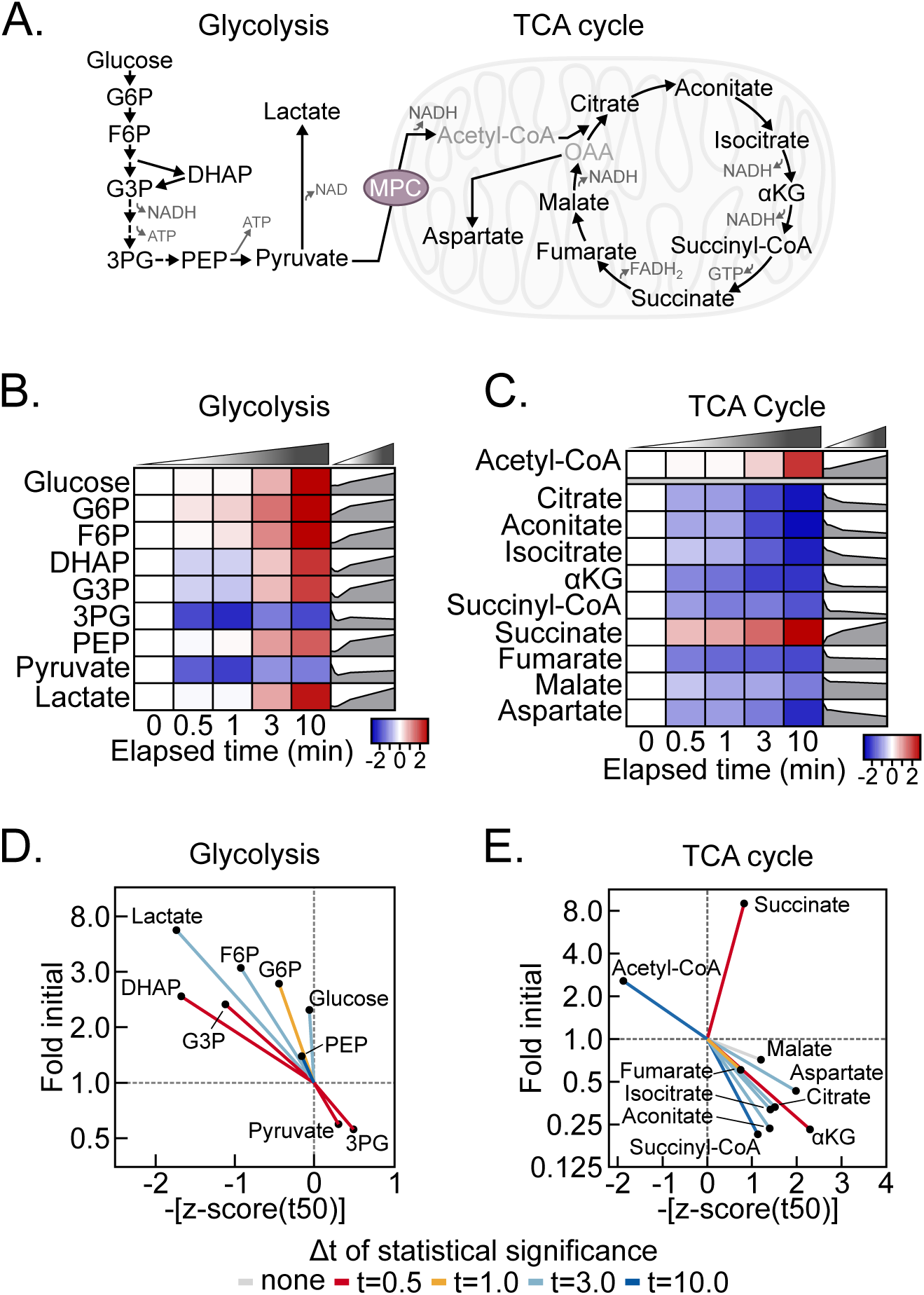
Post-dissection hypoxia rapidly alters central carbon metabolism and purine degradation. (A) Schematic of glycolysis and the TCA cycle. G6P, glucose 6-phosphate; F6P, fructose 6-phosphate; DHAP, dihydroxyacetone phosphate; G3P, glyceraldehyde 3-phosphate; 3PG, 3-phosphoglycerate; PEP, phosphoenolpyruvate; αKG, α-ketoglutarate; OAA, oxaloacetate; MPC, mitochondrial pyruvate carrier. (B-C) Heatmaps depicting relative change of the glycolytic metabolites (B) and TCA cycle metabolites (C) at the indicated time versus t=0. Data are presented as the mean/time point. The simple line graphs were generated using fold initial data calculated relative to t=0. (n=5-6 biological replicates) (D-E) Urchin plots of the glycolytic metabolites (D) and TCA cycle metabolites (E). Lines are colored as described in Figure 1D. (n=5-6 biological replicates) Related to Figure S2.

Urchin plots for both glycolysis and the TCA cycle further revealed distinctive features of post-dissection, ex vivo metabolism (Figure 3D, 3E). For glycolysis, increased versus decreased metabolites binned oppositely to the upper left and lower right quadrants, respectively. Because glucose also increased, this is consistent with a relatively sustained rate of glycogenolysis feeding the overall increases in glycolytic intermediates. In contrast to glycolysis, TCA cycle metabolites strongly binned to the lower right quadrant, with succinate markedly diverging in the upper right (Figure 3E). This suggests that TCA cycle metabolites are highly sensitive to hypoxia and is consist with bi-directional flux into succinate as a metabolic dead-end during hypoxia. In addition to succinate, acetyl-CoA binned distinctly, to the upper left, possibly because of cytosolic accumulation. Together, these data demonstrate complex, time-dependent responses of glycolysis and the TCA cycle to dissection-induced hypoxia. This is important to consider for rigorous metabolomic analysis, because it demonstrates that metabolomic changes within and across adjoining pathways during delayed freezing are not equal, and thus they do normalize away during data analysis.

### Post-dissection liver hypoxia leads to enormous increases in purine degradation metabolites

To identify high dynamic-range markers of post-liver dissection, hypoxia-induced metabolomic drift, we looked for relationships among the metabolites with the highest fold increases (Figure 4A). Strikingly, the six most increased metabolites were products of the redox- and oxygen-regulated purine degradation pathway (Figure 4B). An urchin plot of the purine degradation pathway shows that its metabolites have distinctive kinetic and magnitude of change features (Figure 4C). In contrast to components of glycolysis and the TCA cycle, all purine nucleotide degradation products increased. Moreover, the line vectors for these metabolites generally centered around the mean line and displayed steep slopes with enormous fold changes. Several purine nucleotide degradation products showed greater than 50-fold increases by 10 minutes, which for inosine and xanthine exceeded 100-fold (Figure 4D-F). This combination of high-magnitude increases and generally near-mean t50 kinetics suggests that purine nucleotide degradation may serve as a representative, sensitive marker of post liver dissection metabolomic decay.

**Figure 4:**
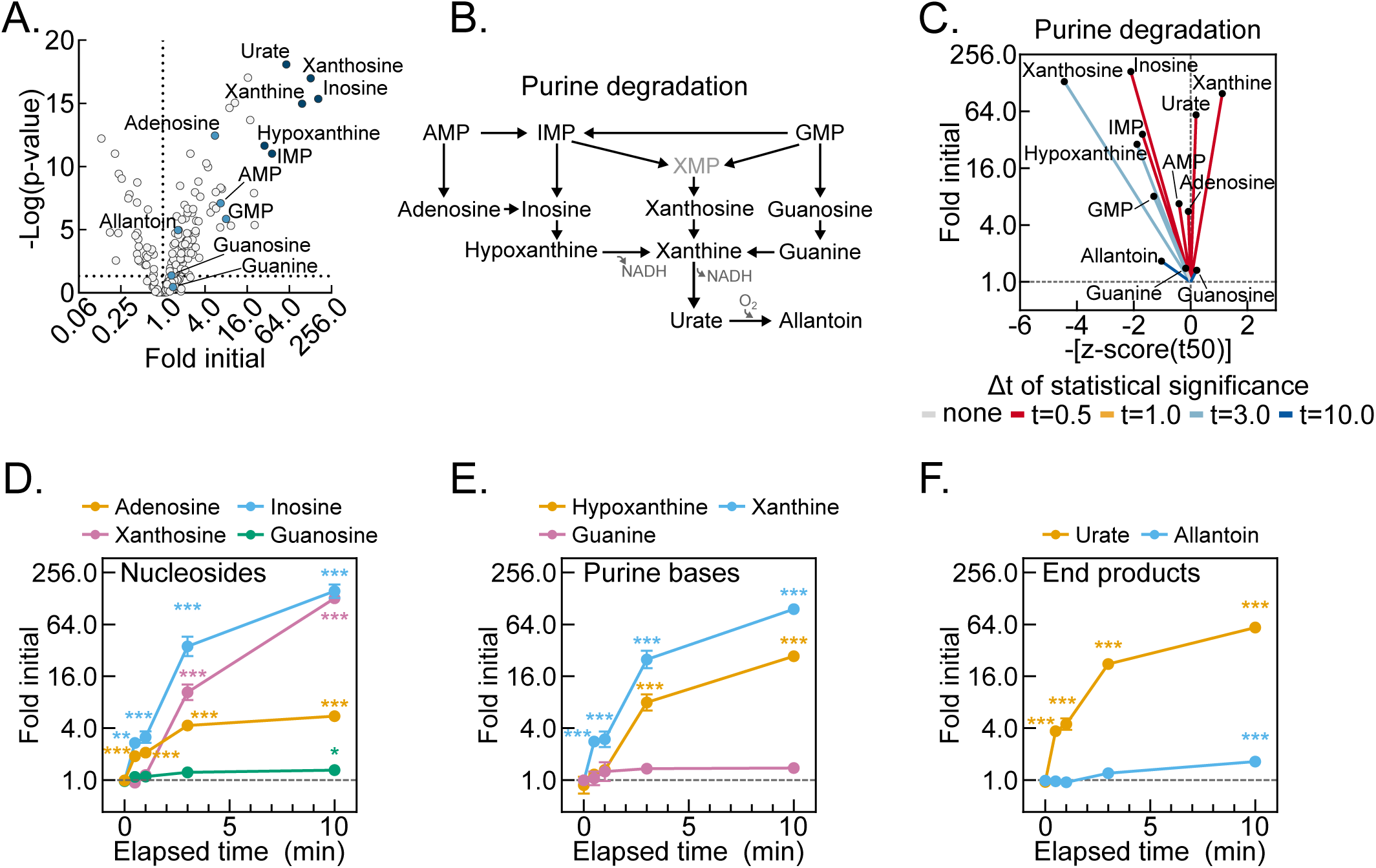
Post-dissection hypoxia rapidly alters purine degradation. (A) Volcano plot of the t=10 data shown in Figure 1C. Metabolites in the purine degradation pathway are and labeled and colored blue. The six greatest fold increased purine degradation products are colored dark blue All other metabolites are colored light gray. (n=5-6 biological replicates) (B) Schematic of the purine degradation pathway. (C) Urchin plot of metabolites in the purine degradation pathway. (n=5-6 biological replicates) (D-F) Levels of purine nucleosides (D), purine bases (E), and purine degradation end products in the liver during the post-dissection time course. (F). Data are presented as the mean±SEM. Statistical significance was calculated using one-way ANOVA with post-hoc Holm-Sidak multiple comparison test versus t=0 on log transformed data. (n=6 biological replicates) *p<0.05, **p<0.01, ***p<0.001.

### Delayed post-dissection freezing of heart and skeletal muscle results in a shift to hypoxic metabolism

Compared to the liver, other tissues may have different metabolomic sensitivities to and signatures of delayed post-dissection freezing. To address this question, we performed test of concept experiments with heart and skeletal muscle, two mouse tissues frequently investigated in the translational metabolism literature (13–15). Heart muscle, like the liver, has a constitutively high metabolic rate but, unlike the liver, its primary function is contractile. Skeletal muscle, like the heart, is striated and has a primary contractile function but, unlike the heart, has a low resting metabolic rate. Thus, comparing metabolomic changes among liver, heart muscle, and skeletal muscle after delayed freezing could inform both generalized properties, tissue-specific functional, and metabolic rate correlates of delayed freezing.

Given the high metabolic rate of heart muscle, we chose a 2-minute freezing delay as a period likely to reveal clear signatures of post-dissection, ex vivo metabolism. Metabolomic change with a 2-minute delay to freeze was substantial. Like the liver, this included many-fold increases in several purine nucleotide degradation products and increases in the NADH:NAD+ and AMP:ATP ratios (Figures 5A, S4A-B, Table S3). Acyl-carnitines, which had increased a few-fold in the liver by delaying freezing 10 minutes, were increased several-fold in the heart at the 2-minute timepoint (Figure 5B, Tables S1, S3). This trend is consistent with the rapid failure of high, baseline β-oxidation. It suggests that the accumulation of acyl-carnitines may be an additional, heart-specific signature of hypoxic metabolism.

**Figure 5:**
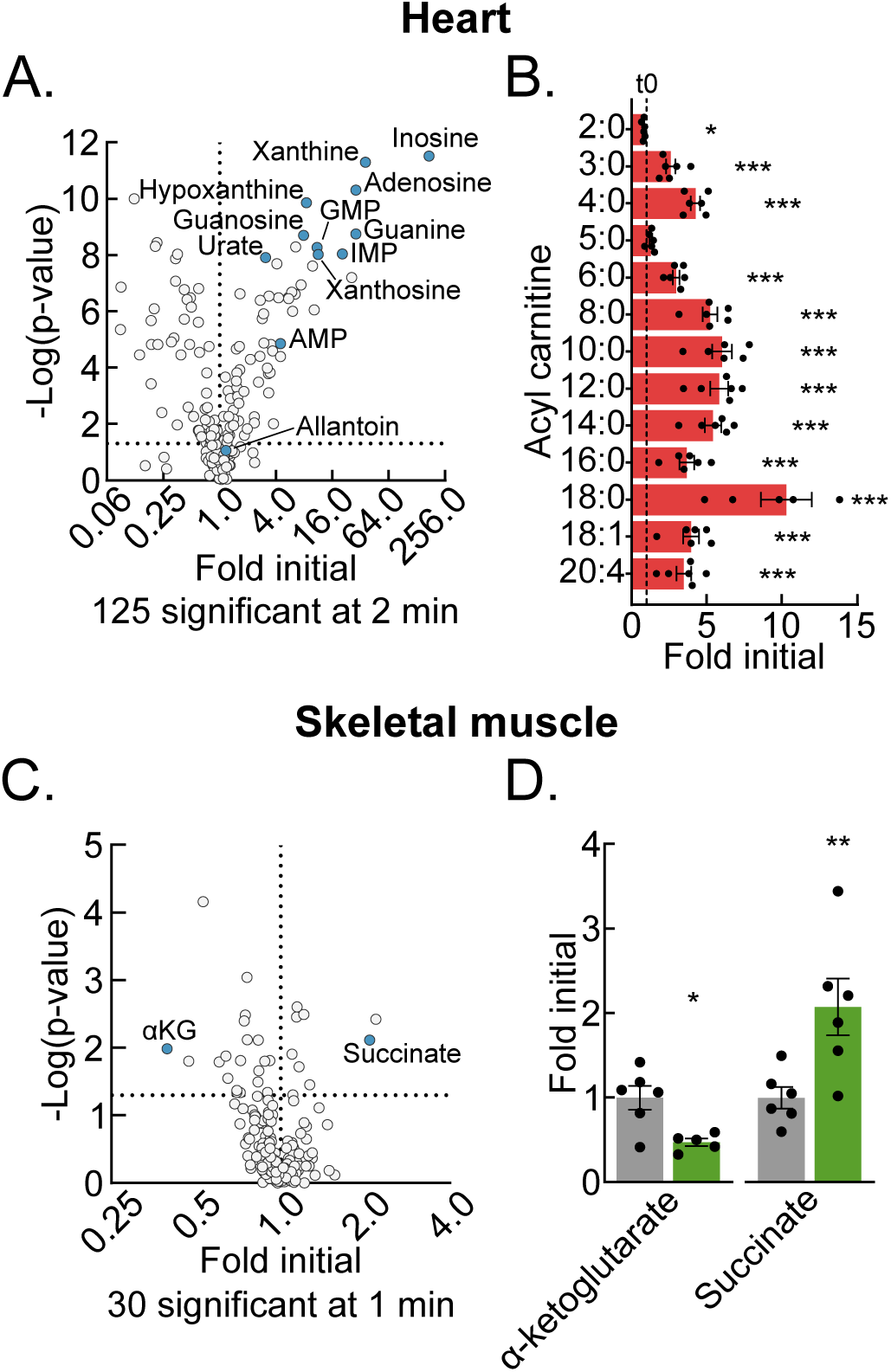
Post-dissection hypoxia rapidly alters the heart and skeletal muscle metabolomes. (A) Volcano plot of the t=2 heart metabolomic profile data compared to t=0. Purine degradation pathway metabolites are colored blue and are labeled. Transformed p-values were calculated by one way ANOVA with post-hoc Holm-Sidak multiple comparison test versus t=0 on log transformed data. (n=5-6 biological replicates) (B) Acyl-carnitine levels for 2-minute heart samples relative to t=0. The dotted line represents the t=0 level (1.0). Data are presented as the mean±SEM. Statistical significance was calculated using the Student’s t-test versus t=0 on log transformed data. (n=5-6 biological replicates) (C) Volcano plot of the t=1 skeletal muscle metabolomic profile data compared to t0. α-ketoglutarate and succinate are colored blue and are labeled. Transformed p-value were calculated using the Student’s t-test versus t=0 on log transformed data. (n=5-6 biological replicates) (D) α-ketoglutarate and succinate levels from t=0 (gray) and t=1 (green) minute skeletal muscle samples. Data presented as the mean±SEM. Statistical significance as calculated using the Student’s t-test. (n=5-6 biological replicates) *p<0.05, **p<0.01, ***p<0.001; Related to Figure S3 and Tables S3 and S4.

Given the slow metabolic rate of resting skeletal muscle, we used a shorter freezing delay to test for a window of metabolomic stability, before the onset of dissection-induced hypoxic metabolism. We utilized the mouse tibialis anterior muscle, which can be quickly dissected to obtain a representative time 0 samples as a control. Thirty metabolites significantly changed with a 1-minute delay to freeze (Figure 5C, Table S4). Notably, the NADH:NAD+ ratio, but not AMP:ATP ratio was significantly changed (Figure S4C-D). Lack of change in the latter may reflect initial buffering by the large skeletal muscle creatine phosphate pool, which decreased but did not reach statistical significance by 1 minute. Like for the liver and heart muscle, the highly oxygen-sensitive TCA cycle metabolite pair, α-ketoglutarate and succinate, which are known to reciprocally regulate several hypoxia-responsive transcriptional control programs, significantly decreased and increased, respectively (Figure 5D). These results from skeletal muscle demonstrate that even metabolically slower tissues must be rapidly frozen after dissection to preserve the in vivo metabolome. Together, data from liver, heart muscle, and skeletal muscle show that although common metabolomic features of hypoxia persist across different tissues, that individual tissues also have unique signatures of post-dissection, hypoxic metabolism.

### Delayed liver freezing leads to both false negative and false positive between-genotype metabolite level differences

A critical question for rigorous metabolomic investigation is how individual metabolite levels in control versus experimentally modified tissues change during post-harvest delays to freeze. If such metabolomic drift across different conditions is similar, then equally delaying freezing time could enable accurate comparisons when immediate freezing is not ideal for experimental design. To test this possibility, we returned to our well-vetted liver system and compared dissection-induced, hypoxic metabolomic drift between WT and liver-specific mitochondrial pyruvate carrier (MPC) knockout (MPC-LivKO) mouse livers. Liver tissue was frozen immediately after 0 (about 1 second) and 1- and 3-minute delays. When frozen immediately, as previously reported, MPC-LivKO and WT livers displayed many between-genotype metabolomic differences (Figure S5A, Table S5) (16–18). To directly compare relative rates of change between individual metabolites, MPC-LivKO and WT metabolite level ratios at all time points were normalized to the time 0 ratio. This normalized the time 0 ratios to 1, with the per metabolite ratios at 1- and 3-minute freezing delays being greater or less than 1 if they increased more or less in MPC-LivKO mice, respectively (Table S6).

Rendering these between-genotype metabolite ratios on a heat map and volcano plot revealed myriad genotypic differences in post-harvest metabolomic change (Figure 6A, S5B). Specific examples of metabolites for which the between-genotype relative abundance changed are: pyruvate (greater initial level in MPC-LivKO was lost); α-ketoglutarate (lesser initial level in MPC-LivKO was lost); acetoacetyl-CoA (similar initial level, a greater MPC-LivKO level was gained); and propionate (greater initial level in MPC-LivKO was inverted to a lesser level) (Figures 6B-E). Thus, genotypic differences in ex vivo tissue metabolism cannot be controlled for by equally delaying time to post-harvest freeze, which leads to both false negative and false positive between-genotype differences.

**Figure 6:**
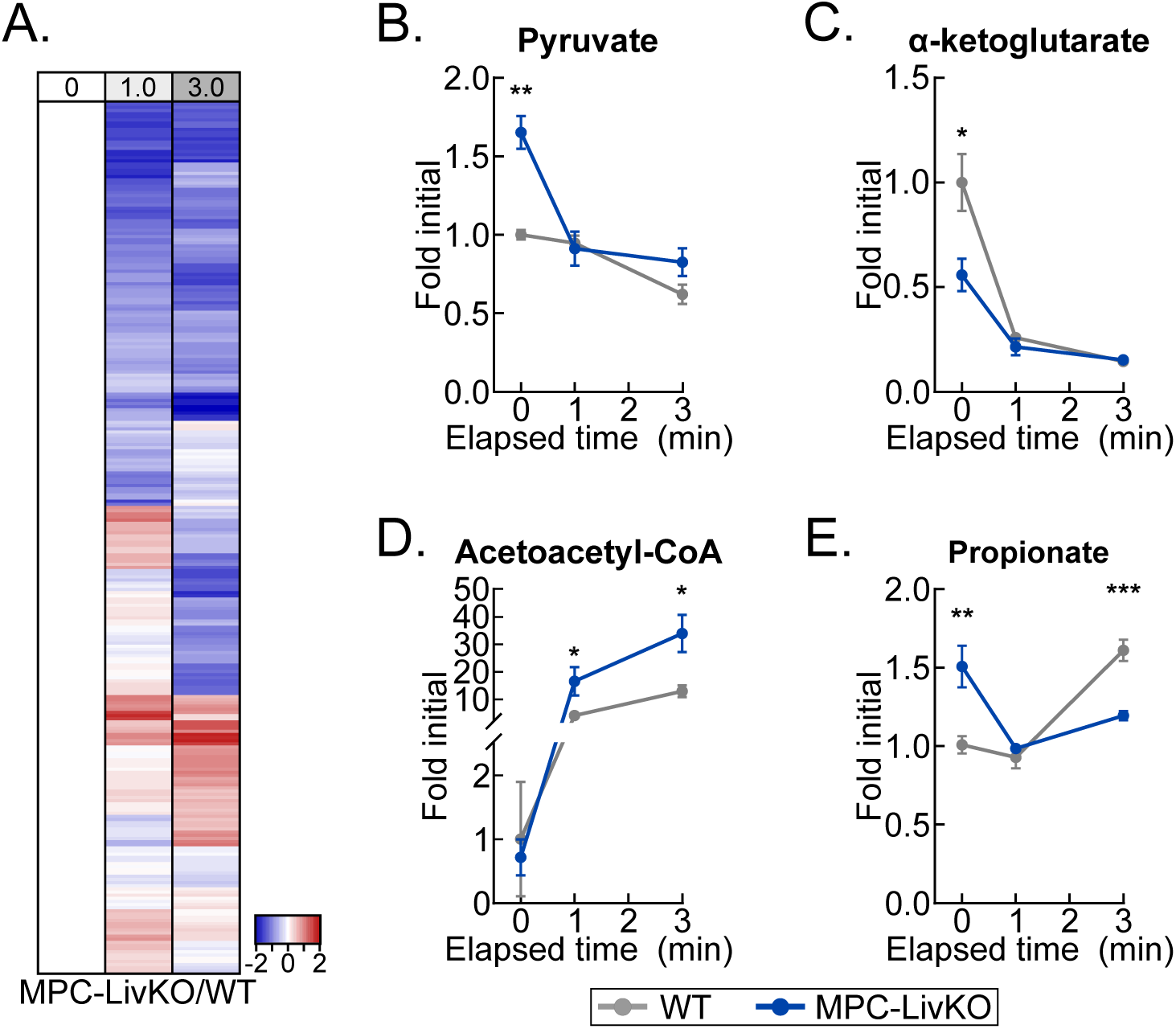
Post-dissection hypoxia rapidly affects phenotypic comparisons and ^13^C-isotopologue enrichments. (A) Heatmap depicting the relative change of the MPC-LivKO/WT liver metabolome throughout the delayed freezing time course versus MPC-LivKO/WT t=0. Data are presented as the mean/time point. (n=4-5 biological replicates) (B-E) Levels of pyruvate (B), α-ketoglutarate (C), acetoacetyl-CoA (D) and propionate (E) in MPC-LivKO (blue) versus WT (gray) livers at t=0. Data are presented as the mean±SEM. Statistical significance for difference between genotypes at each time was calculated using the Student’s t-test on log transformed data. (n=4-5 biological replicates) *p<0.05, **p<0.01, ***p<0.001; Related to Figure S4 and Tables S5, and S6.

### Post-dissection, ex vivo liver metabolism changes ^13^C isotopologue abundances and fractional enrichments

Finally, we asked whether liver ^13^C isotopologue enrichments following [U^13^C]-glucose administration are subject to post-harvest change. Stable-isotope tracing is increasingly being employed to investigate substrate-specific metabolism over minutes- to hours-long time intervals (19, 20). This raises the possibility that such a time-integrated signal is more stable than the broad metabolome during post-dissection hypoxia. We administered a single bolus injection of [U-^13^C]-glucose and examined changes in liver lactate and TCA cycle ^13^C-enrichments, following 1- and 3-minute freezing delays (Figures 7A, Table S7). We observed striking post-harvest changes in both metabolite pool sizes and percent ^13^C enrichments (Figures 7B-G). Notably, the percentage of the succinate m+3 isotopologue increased many-fold (Figure 7E), which is consistent with increased flux through pyruvate carboxylation and TCA cycle reversal (Figure 7A). Together, these data show that, like metabolite levels, isotopologue abundances and enrichment percentages change with post-dissection freezing delays. This likely represents a shift in the proportional flux from metabolite feeder pools. It demonstrates the requirement to rapidly freeze tissues after dissection to obtain ^13^C tracing data reflecting the in vivo state.

**Figure 7:**
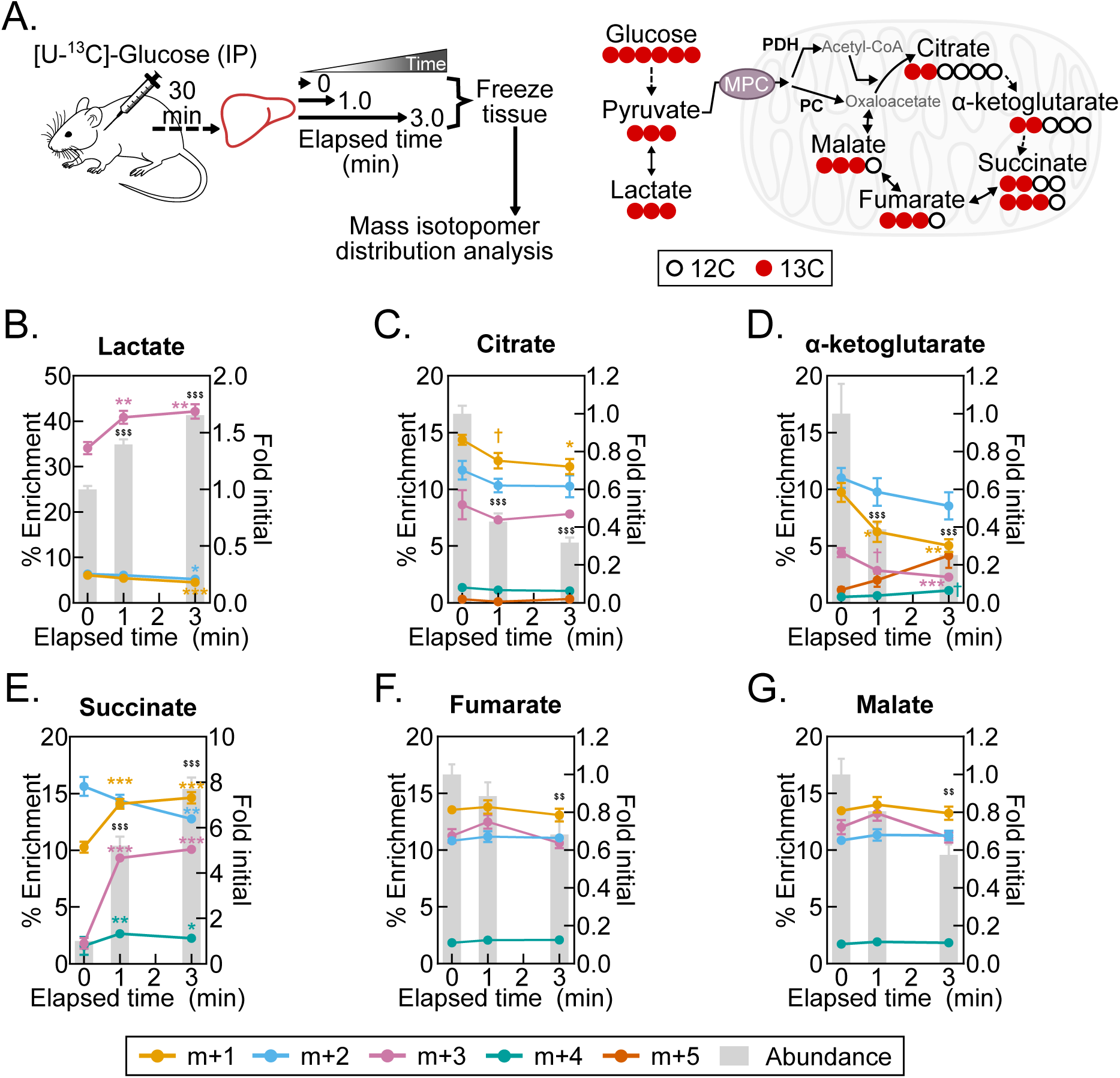
Post-dissection hypoxia rapidly affects ^13^C-isotopologue enrichments. (A) Schematics of experimental procedures for [U-^13^C]-glucose tracing during hypoxia induced by tissue-dissection and ^13^C incorporation into the TCA cycle following administration of [U-^13^C]-glucose. (B-G) Overlays of % isotopologue enrichment line graphs (left y-axis) with fold initial abundance bar graphs (right y-axis) for lactate (B), citrate (C), α-ketoglutarate (D), succinate (E), and fumarate (F), and malate (G). Data are presented as the mean±SEM. Statistical significance of difference was calculated using one-way ANOVA with post-hoc Holm-Sidak multiple comparison test versus t=0 on log transformed data. ^$$^p<0.01 and ^$$$^p<0.001 for fold initial abundance bar graphs. (n=8 biological replicates) ^†^p<0.1, *p<0.05, **p<0.01, ***p<0.001 (% Enrichment); ^$$^p<0.01, ^$$$^p<0.001 (Fold initial); Related to Tables S6 and S7.

## Discussion

Metabolomics is a powerful, mainstreaming research approach that is frequently applied to animal tissues to understand in vivo metabolism. Because broad metabolomic analysis of tissues requires dissection, a major risk to the integrity of metabolomic data is a hypoxia-induced shift of tissue metabolism between the time of disrupted tissue perfusion and freezing. However, most manuscripts reporting metabolomic analysis of mouse or other tissues do not precisely specify the lag time between loss of perfusion and tissue freezing. Thus, it has remained unclear how delayed tissue freezing affects multiple classes of tissue metabolomic data. We addressed this problem by carefully and systematically assessing broad, genotype-specific, and ^13^C isotopologue metabolomic change resulting from post-dissection, ex vivo mouse tissue metabolism.

Prior work addressing metabolomic change with delayed tissue freezing has been limited in scope or without seconds-to-single minutes coverage of the immediate post-dissection period (21–23). Notably, the most recent of these carefully demonstrates that delaying mouse liver freezing by as little as 10 seconds alters the lactate:pyruvate ratio and TCA cycle intermediate levels (23). Although the implications of these findings are critical to the field, how they extend to the larger metabolome has remained unaddressed. Our findings here corroborate this observation and show how it fits within a framework of broad, interdependent post-dissection metabolomic drift. One NMR-based study of intact tumor pieces reports that 16 metabolites in mouse tumor xenografts changed only minimally after 30 minutes but significantly after 90 minutes ex vivo, after removal (24). However, this publication does not specify the time that elapsed between loss of perfusion and the time 0 reference measurement. Thus, substantial metabolomic drift could have occurred and minimized observable changes by the 30-minute time point. Conversely, the observed changes after the very long delay of 90 minutes might have been chemical versus enzymatic, obscuring the possibility that enzymatic metabolism had ceased much earlier in the time course. In a study of metabolomic and macromolecular change in primary human tumors, only 10% of numerous measured metabolites, lipids, and peptides were found to be affected by a 3-hour delay in freezing, leading to the conclusion that no specific pathway-wide changes occurred. However, the tumors were cut into pieces prior to freezing and, as in the above study, the lag time from loss of perfusion to freezing of the pieces was not stated (25). Although incomplete technical detail occludes clear interpretation of the data, it is also possible that the findings of lower sensitivity of tumors to hypoxia reflect prior biological adaptation to a lack of perfusion in vivo (26–28). Future, carefully controlled studies will be required to determine this, which may also apply to poorly vascularized normal tissues like cartilage or other connective tissues. Thus, in comparison to these prior studies, our findings here show that delayed freezing after dissection of liver, heart muscle, and skeletal muscle leads to rapid, extensive metabolomic drift.

In addition to evaluating the broad metabolome, here we investigated how tissue ^13^C mass isotopologues change with delayed freezing. In a test of concept experiment, we observed that delayed freezing of the liver leads to altered TCA cycle intermediate ^13^C isotopologue abundances and fractional enrichments, which was especially prominent for succinate. Numerous prior studies have shown that succinate accumulates with ischemia, but they have not addressed how ex vivo ischemic metabolism impacts ^13^C labeling (29–31). Our observation that delayed freezing of liver of mice administered ^13^C glucose results in not only an increase in the size of the succinate pool but also a many-fold increase in the m+3 succinate ^13^C isotopologue is consistent with reverse TCA cycle flux driven by a high NADH:NAD+ ratio. It also suggests that isotopologue distributions of metabolites with differentially redox-regulated feeder pools may be especially sensitive to delays in tissue freezing.

Given that ex vivo tissue metabolism after dissection rapidly alters the broad in vivo and isotope-enriched metabolome, we highlight best practices for sample collection to obtain accurate in vivo metabolomic data. For most tissues, the need for rapid, controlled freezing can be accommodated by using a liquid nitrogen-cooled Wollenberger-like freeze-clamp device. This instrument maximizes the heat transfer rate and results in nearly instantaneous sample freezing (32). Therefore, samples destined for metabolomics must be prioritized, and it is likely not possible to preserve the in vivo metabolome of multiple tissues collected from a single animal. Here, the liver and heart showed rapid, high magnitude increases in purine nucleotide degradation products that could serve as a within-study marker of metabolomic quality for these tissues. This is also consistent with the elevated purine degradation products observed in the plasma of hypoxic newborn humans, mouse retinas after extended incubation in culture media, and mouse tissues with protracted ischemia (22, 31, 33). Although such changes were not observed in skeletal muscle following a 1-minute delay in freezing, the α-ketoglutarate:succinate ratio decreased several-fold, suggesting this ratio could serve as a skeletal muscle-specific marker of post-dissection, ex vivo metabolism. Thus, when investigating animal tissue metabolomics, tissue- and condition-specific pilot experiments to determine signatures of metabolomic change after loss of perfusion may be useful for informing data quality.

Lastly, we note key limitations of this study, which was not designed to investigate variables other than how mouse tissue dissection-induced hypoxia alters the in vivo metabolome. First, we do not account for the effects of anesthesia, which has previously been shown to impact the metabolome (34, 35). For purposes of experimental control, we used isoflurane as a rapidly acting anesthesia with predictable dose- and timing-dependent effects. However, the time 0 values that we calculate fold changes from could differ by anesthetic, which could influence fold-change results. Second, compared to humans and larger animals, mice have a much higher metabolic rate. Thus, the rate of post-dissection metabolomic drift in humans and other animals may be slower, which may permit mild freezing delays before the onset of confounding metabolomic drift. Nonetheless this possibility needs to be experimentally tested. And third, although we demonstrate that post-dissection metabolomic change is a major risk to research rigor and data integrity, we speculate that such change nonetheless may inform biology. As we observed comparing WT MPC and LivKO liver tissue, delayed freezing led to both false negative and false positive between-genotype differences. Because these differential, genotype-specific rates of metabolomic change reflect differences in metabolic programming, they may inform secondary mechanistic experiments to define the underlying molecular bases. Yet, even in this case, to be most useful, a well-controlled time zero is still required to fully detect such ex vivo metabolomic divergence.

In conclusion, in this study we illustrate that delayed mouse tissue freezing leads to rapid, extensive metabolomic remodeling. This is consistent with the fact that, compared to the proteome and transcriptome, the metabolome is highly dynamic, responding nearly instantaneously to metabolic stimuli and other cues. Whereas the transcriptome and base proteome, independent of modification state, are stable for minutes to hours after tissue dissection, we show here the that the broad metabolome of three mouse tissues markedly changes in the early seconds and minutes after dissection. Because obtaining accurate results requires that the in vivo metabolome is fixed prior to analysis, achieving research rigor in metabolomics brings unique and unavoidable challenges for accurate experimental design. These additional challenges to ensuring data integrity, combined with the rapidly escalating use of metabolomics as an investigative method, make this an important time for the field to embrace best practices. We suggest that all manuscripts reporting metabolomic data provide clear, precise information about the time that elapsed between loss of tissue perfusion and tissue freezing, as well as minimizing these times as much as possible.

## Supporting information

Supplemental Object1

## Acknowledgements

We are grateful to the University of Iowa Fraternal Order of Eagles Diabetes Research Center Metabolomics Core Facility for technical assistance. This work was supported by grants NIH R01 DK104998 (EBT), University of Iowa Healthcare Distinguished Scholars Award (EBT), T32 DK112751 to E. Dale Abel (DAS), ADA 1-18-PDF-060 and AHA CDA851976 (AJR), and T32 GM007337 to Steven Lentz (DJP). We are grateful to Dr. Bo Ram Kim, Dr. Alejandro Pezzulo, and Dr. Leonid Zingman, and the Scientific Editing and Research Communication Core (Dr. Christine Blaumueller), at the University of Iowa Carver College of Medicine for critically evaluating a draft manuscript.

## Author Contributions

EBT conceived the project. AJR and EBT designed experiments. AJR, DJP, ASK, and DAS performed experiments. EBT, AJR, and NB performed data analysis and data visualization. AJR, NB, and EBT wrote and edited the manuscript. All authors read and commented on a draft manuscript.

## Supplemental object 1

Interactive and fully zoomable version of the urchin plot shown in Figure 1D. Urchin plot produced using Plotly. Place your cursor over the terminus of a line to view its label.

Table S1: Metabolomics data for liver hypoxia time course, for individual samples

Table S2: Urchin plot t50 calculations from liver data

Table S3: Metabolomics data for heart hypoxia at t=0 and t=2, for individual samples

Table S4: Metabolomics data for skeletal muscle hypoxia at t=0 and t=1, for individual samples

Table S5: Liver metabolomics data for MPC-LivKO versus WT at t=0, t=1, and t=3, for individual samples

Table S6: Liver metabolomics data for MPC-LivKO/WT at t=1 and t=3 relative to MPC-LivKO/WT t=0, for individual samples

Table S7: Liver U-^13^C-glucose isotopologue enrichment data for individual samples

## Resource availability

### Lead contact

Additional information and requests for reagents and resources should be directed to and will be fulfilled by the lead contact, Dr. Eric Taylor.

### Materials availability

No new materials were generated during this study.

## Materials and methods

### Materials

C57bl6/j mice were obtained from Jackson Laboratories (Stock #000664). Internal standards citric acid (2,2,4,4-D4, 98%; #DLM-3487), succinic acid (2,2,3,3-D4, 98%; #DLM-584), L-valine (2,3,4,4,4,5,5,5-D8; 98%; #DLM-311), L-glutamic acid (^13^C5, 99%; #CLM-1800), L-glutamine (^13^C5, 99%, #CLM-1822), L-lysine (^13^C6, 99%; #CLM-2247), L-methionine (^13^C5, 99%; #CLM-893), and L-tryptophan (^13^C11, 99%; #CLM-4290) and tracer glucose (^13^C6, 99%, #CLM-1396) were obtained from Cambridge Isotope Laboratories, Inc. AAV8-TBG-NULL (#105536-AAV8) and AAV8-TBG-Cre (#107787-AAV8) were purchased from Addgene. Pre-filled bead mill tubes containing 1.4 mm ceramic beads (#15-340-153) were purchased from FisherScientific. Methoxaymine hydrochloride (#226904, MOX) and N-methyl-N-(trimethylsilyl) trifluoroacetamide (#694709, MSTFA) were purchased from Sigma Aldrich. Protease inhibitor cocktail (#786-437) was purchased from G Biosciences. Bio-Rad protein assay dye regent (#5000006) and 0.45 µm nitrocellulose membrane (#1620115) were purchased from Bio-Rad.

### Animal sacrifice

All animal work was performed in accordance with the University of Iowa Animal Use and Care Committee (IACUC) guidelines. For all studies, mice were fasted 4 hours before sacrifice. 9–11-week-old male mice were used for WT liver, TA and U-^13^C-glucose tracing studies. 15-week-old male mice were used for the WT and MPC-LivKO liver study. Mice were anesthetized by 3.5% isoflurane inhalation for 1.75 minutes prior to tissue dissection. Following dissection, tissue was frozen immediately (t=0) or maintained in a humidified chamber for up to 10 minutes before freezing (t=10). A liquid-nitrogen cooled Wollenberger-like device referred to here as a freeze-clamp was used to freeze tissue samples instantaneously.

In the case of liver samples, the thoracic cavity was opened to expose the liver, and care was taken not to disrupt or puncture the diaphragm. The left lateral lobe of the liver was isolated and dissected from anesthetized mice. The freeze-clamp was used to freeze liver samples at the appropriate time points.

For heart tissue samples: the thoracic cavity was opened to expose the diaphragm. At the time of collection, the diaphragm and sternum were quickly cut to open the pericardial space and expose the heart. Forceps were used to quickly isolate the heart, and ventricular tissue was separated from the atria. Ventricular tissue was frozen at the appropriate time point by using te freeze clamp. Freeze-clamping the cardiac tissue as described resulted in the expulsion of most ventricular blood.

For skeletal muscle samples: the skin was carefully removed from the leg of the anesthetized mouse. The tibialis anterior muscle (TA) was identified, and the fascia was removed. The fine tip of a micro-fine tweezer was placed under the TA muscle without disrupting the attached tendons. At the time of dissection, the micro-fine tweezer tip was used to disrupt one TA muscle tendons and the second tendon was cut, thereby liberating the TA muscle, which was frozen at the appropriate time point using the freeze-clamp.

### Generation of Mpc1 liver-specific knockout mice

Mpc1 liver specific knockout (MPC-LivKO) and littermate paired control (WT) mice were generated as previously reported (16). Briefly, 8-week-old male Mpc1-floxed mice were retroorbitally injected with 1x10^11^ gc/mouse of either AAV8-TBG-NULL or AAV8-TBG-Cre generating WT and MPC-LivKO mice, respectively.

### U-^13^C-glucose tracing

C57Bl6j mice were i.p. injected with U-^13^C-glucose at a dosage of 1.4 g/kg body mass. The mice were anesthetized 30 minutes after injection, as described above, and liver tissue was harvested.

### Western Blots

Freeze-clamped liver samples were homogenized in 40 volumes:weight of a buffer containing 40mM HEPES, 120mM NaCl, 50mM NaF, 5mM sodium pyrophosphate decahydrate, 5mM b-glycerolphosphate, 1mM EDTA, 1mM EGTA, 10% glycerol (v/v), 1% igepal CA-630 (v/v), with 1X protease inhibitor and 1μM DTT. Homogenates were rotated at 4°C for 30 minutes and centrifuged at 21,000 x g. Supernatants were collected, and protein concentrations were determined using the Bio-Rad protein assay reagent. Proteins were separated using TRIS-SDS-PAGE gel electrophoresis, transferred to 0.45 μM nitrocellulose membranes, and blocked with TBST (50mM Tris, 150mM NaCl, and 0.05% Tween-20) supplemented with 2.5% BSA. Blocked membranes were incubated with primary antibodies at 4°C overnight [mouse monoclonal anti-AMPKα (F6) (RRID: AB_915794), rabbit monoclonal anti-pAMPKα (Thr172) (40H9) (RRID: AB_331250), and rabbit monoclonal anti-HSP90 (RRID: AB_2121214) from Cell Signaling Technology]. The following day membranes were washed with TBST and incubated with fluorescently labeled secondary antibodies for 1 hour at room temperature [goat anti-Mouse Dylight 800 (RRID: AB_2556756), donkey anti-Rabbit DyLight 680 (RRID: AB_2556622), and donkey anti-Rabbit DyLight 800 (RRID: AB_2556616) from ThermoFisher]. Membranes were visualized using the Li-Cor Odyssey CLx system (Li-Cor Biosciences).

### Metabolite extraction and derivatization

Liver tissue (40 ± 5 mg) was weighed without thawing, and tissue was lyophilized overnight. Heart tissue and TA muscles were weighed and lyophilized intact. Tissues were extracted in 18 volumes (relative to wet tissue weight) of ice-cold extraction solvent composed of acetonitrile:methanol:water (2:2:1) containing 1 μL/mL of an internal standard mix (0.33 mg of each/mL in water, see “Materials”). Lyophilized tissue was added to pre-filled bead mill tubes containing 1.4 mm ceramic beads. Ice-cold extraction solvent was added, and tissue was homogenized using a BeadRupter bead mill homogenizer (Omni International) for 30 seconds at 6.45 MHz. Immediately afterwards homogenization tubes were rotated for 60 minutes at -20°C. Samples were centrifuged at 21,000 x *g* for 10 minutes, after which the supernatant was transferred to a new 1.5 mL microcentrifuge tube and pulse vortexed to ensure uniform mixing of the supernatant. Of this supernatant, 150 μL was transferred to glass autosampler vials for GC-MS analysis and 300 μL was transferred to a 1.7 ml microcentrifuge tube for further processing for LC-MS analysis. A quality control (QC) sample was prepared by pooling equal volumes from each sample; the QC sample was aliquoted into a glass autosampler vial and a 1.7 ml microcentrifuge tube for GC-MS (150 μL) and LC-MS analysis (300 μL), respectively. Samples were dried in a Speedvac Vacuum concentrator (Thermo) for 2 hours without heating at a vacuum ramp = 4.

### GC-MS Analysis

GC-MS sample derivatization with MOX+MSTFA was accomplished as follows: 30 μL of pyridine containing 11.4 mg/mL of MOX was added to the dried metabolite extracts. Samples were then vortexed for 10 minutes and heated at 60°C for 60 minutes. Next, 20 uL of MSTFA was added to the pyridine/MOX derivatized samples, and they were vortexed for 5 minutes, and heated at 60°C for an additional 30 minutes. GC-MS analysis was performed using a Trace 1300 GC (Thermo) coupled to an ISQ-single quad mass spectrometer (Thermo). For each sample, 1 μL of sample was injected into the GC by an autosampler in split mode (split ratio: 20-1; split flow: 24 μL/minute, purge flow: 5 mL/minute, Carrier mode: Constant Flow, Carrier flow rate: 1.2 mL/minute). Separation was accomplished using a standard fused silica TraceGold TG-5SilMS column (Thermo). The temperature gradient was as follows: 80°C for 3 minutes, temperate was ramped at a rate of 20°C/minute to a maximum temperature of 280°C and held for 8 minutes.

Between sample runes, the injection syringe was washed 3x with methanol and 3x with pyridine. The MS was operated from 3.90 to 21.00 minutes in EI mode (-70eV) using select ion monitoring (SIM). The instrument was tuned and calibrated daily. The pooled QC sample was analyzed at the beginning and at the end of the GC/MS run, as well as intermittently throughout.

### LC-MS analysis

For LC-MS analysis, dried metabolite extracts were resuspended in 30 ul of acetonitrile:water (1:1), vortexed for 10 minutes, and stored at -20°C overnight. The following day, resuspended samples were centrifuged at 21,000 x *g* for 10 minutes, and the resulting supernatant were transferred to autosampler vials for LC-MS analysis. For each prepared sample, 2 µL were separated using a Millipore SeQuant ZIC-pHILIC (2.1 X 150 mm, 5 µm particle size) column with a ZIC-pHILIC guard column (20 x 2.1 mm) attached to a Thermo Vanquish Flex UHPLC. The mobile phase was comprised of Buffer A [20 mM (NH4)2CO3, 0.1% NH4OH (v/v)] and Buffer B [acetonitrile]. The chromatographic gradient was run at a flow rate of 0.150 mL/min as follows: 0– 21 min-linear gradient from 80 to 20% Buffer B; 20-20.5 min-linear gradient from 20 to 80% Buffer B; and 20.5–28 min-hold at 80% Buffer B. Data was acquired using a Thermo Q Exactive MS operated in polarity switching fullscan mode with a spray voltage set to 3.0 kV, the heated capillary held at 275°C, and the HESI probe held at 350°C. The sheath gas flow was set to 40 units, the auxiliary gas flow was set to 15 units, and the sweep gas flow was set to 1 unit. MS data resolution was set at 70,000, the AGC target at 10e6, and the maximum injection time at 200 ms. The QC sample was analyzed at the beginning and at the end of the LC-MS run, as well as intermittently throughout.

### Mass Spectrometry Data Analysis

Acquired GC- and LC-MS data were processed using the Thermo Scientific TraceFinder 4.1 and TraceFinder 5.1 software. Targeted metabolites were identified based on the University of Iowa Metabolomics Core facility standard-confirmed, in-house library containing target ion and at least 1 confirming ion and retention time (GC) or accurate mass, retention time, and MS/MS data (LC). The NOREVA tool used the QC sample analyzed throughout the instrument run to apply local polynomial fits to metabolite peak areas and correct for instrument drift (36). NOREVA corrected data were normalized to the D4-succinate signal/sample to control for extraction, derivatization (GC), and/or loading effects. For ^13^C-tracing analysis, ^12^C-natural abundance was corrected using previously defined equations (37).

### Quantification and statistical analysis

Post NOREVA corrected, D4-succinate-normalized metabolomic data were log transformed. Following log transformation, the Grubbs’ test was employed at an α=0.01 to detect extreme outliers. This resulted in the exclusion of 130/20388 or 0.64% of values. Metabolomic data were then analyzed using an unpaired Student’s t-tests or a one-way ANOVA with the Holm-Sidak posthoc multiple comparison test versus t = 0. Statistically significance differences were defined as having p-values < 0.05 (*), p < 0.01 (**) and p < 0.001 (***). Results were plotted as mean ± standard error of the means (SEM) relative to t = 0. The number of biologically unique samples used in each experiment is specified in the relevant figure legend.

For calculation of total metabolomic change and t50, normalized metabolite values were processed in R (v4.1.0). For each metabolite, a smooth spline model was fitted for the time course using the npreg (v1.0-7) R package and the maximum likelihood methods for smoothing (38). Maximum likelihood method was selected due to fewest issues with overfitting of splines after comparison with generalized cross-validation, Bayesian information criterion, and Akaike’s information criterion. Time at which 50% of the total change in the metabolite occurred was estimated using the direction and magnitude of the largest change of the metabolite and the fitted smooth spline model. The output of this analysis was plotted using the gpplot2 (3.3.5) R package.

---

**Figure S1:**
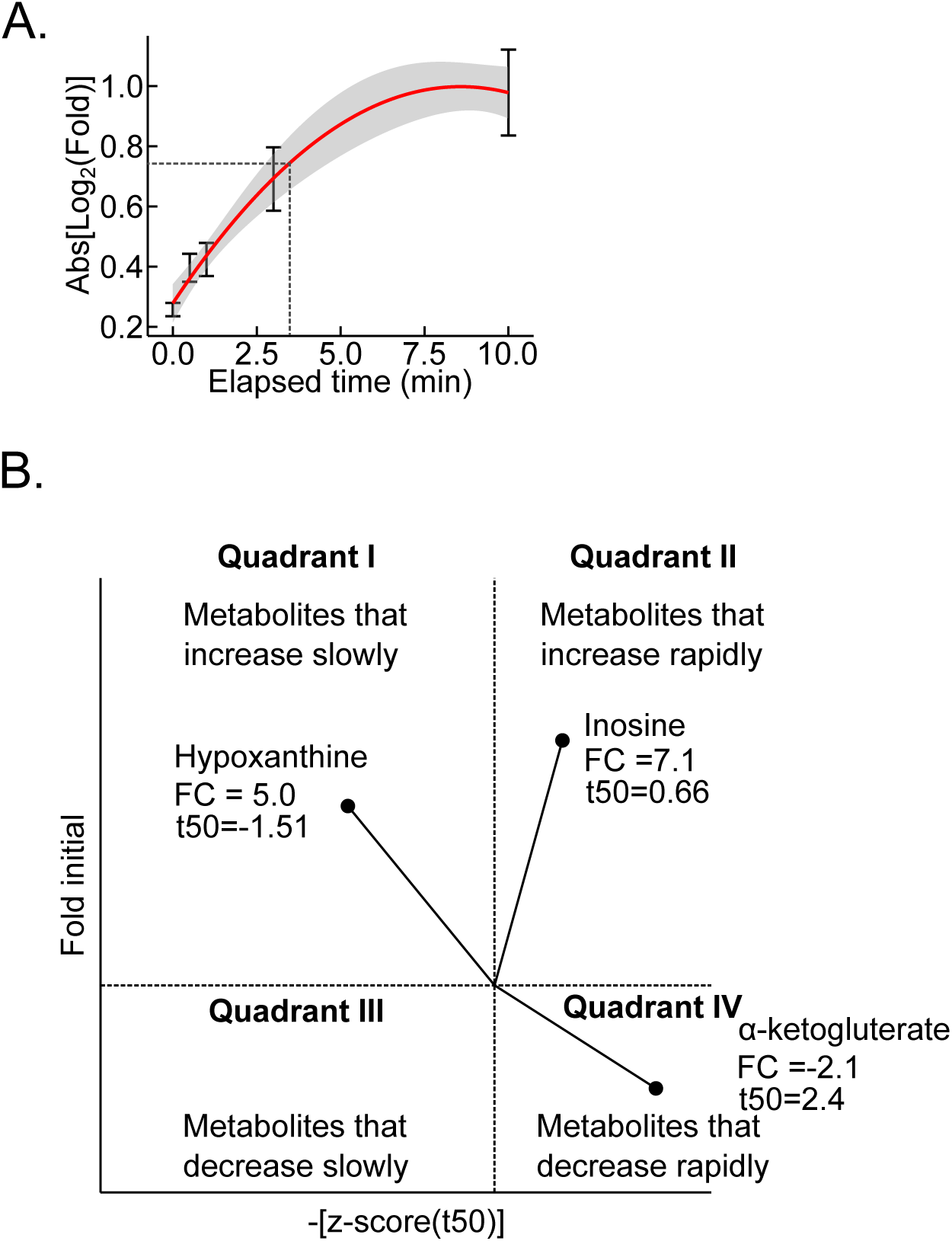
Post-dissection hypoxia results in rapid metabolomic change. (A) Global kinetic change in the measured liver metabolome following tissue dissection. The gray shaded area represents the 95% confidence interval. (n=5-6 biological replicates/metabolite) (B) Example “Urchin plot” with quadrant labels and example metabolites.

**Figure S2:**
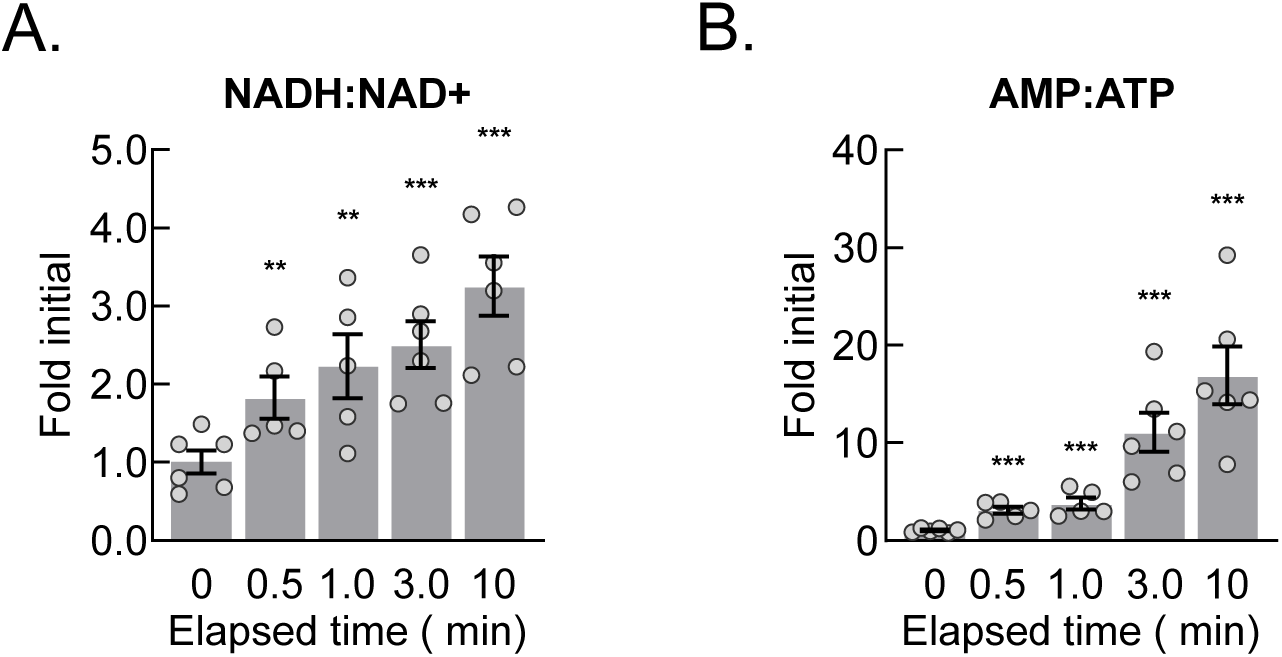
Post-dissection hypoxia alters redox and energetic status. (A-B) The relative liver NADH:NAD+ (A) and AMP:ATP (B) ratios throughout the post-dissection time course. Data are presented as the mean±SEM. Statistic were calculated by one-way ANOVA with post-hoc Holm-Sidak multiple comparison test versus t=0 on log transformed data. (n=5-6 biological replicates) **p<0.01, ***p<0.001.

**Figure S3:**
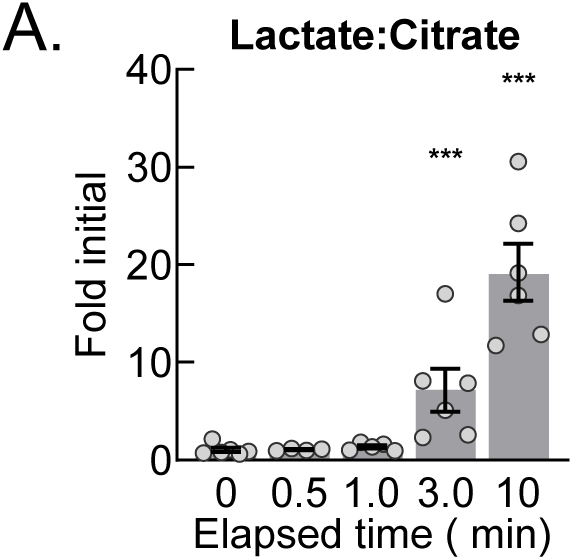
Post-dissection hypoxia results in a switch to anerobic metabolism. (A) The relative lactate:citrate ratio in the liver throughout the post-dissection time course. Data are presented as the mean±SEM. Statistics were calculated using one-way ANOVA with post-hoc Holm-Sidak multiple comparison test versus t=0 on log transformed data. (n=5-6 biological replicates) ***p<0.001.

**Figure S4:**
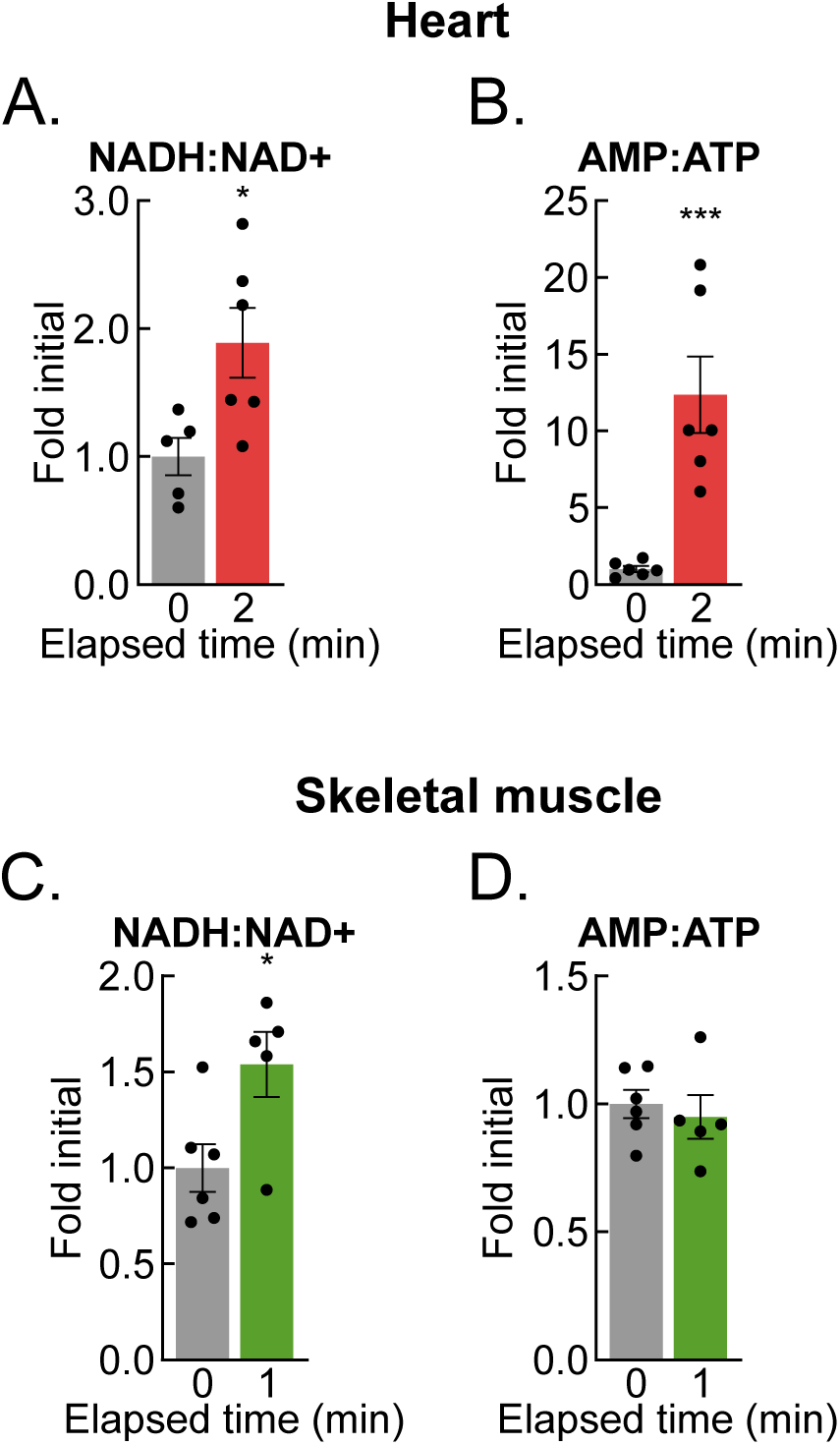
Post-dissection hypoxia results in altered heart and skeletal muscle metabolomes. (A-B) The relative ratios of NADH:NAD+ (A) and AMP:ATP (B) in the heart at t=0 (gray) and t=2 (red) minutes. Data are presented as the mean±SEM. Statistical significance was calculated using the Student’s t-test on log transformed data. (n=5-6 biological replicates) (C-D) The relative ratios of NADH:NAD+ (C) and AMP:ATP (D) in skeletal muscle at t=0 (gray) and t=1 (green) minutes. Data are presented as the mean±SEM. Statistical significance calculated using the Student’s t-test on log transformed data. (n=4-6 biological replicates) *p<0.05, ***p<0.001.

**Figure S5:**
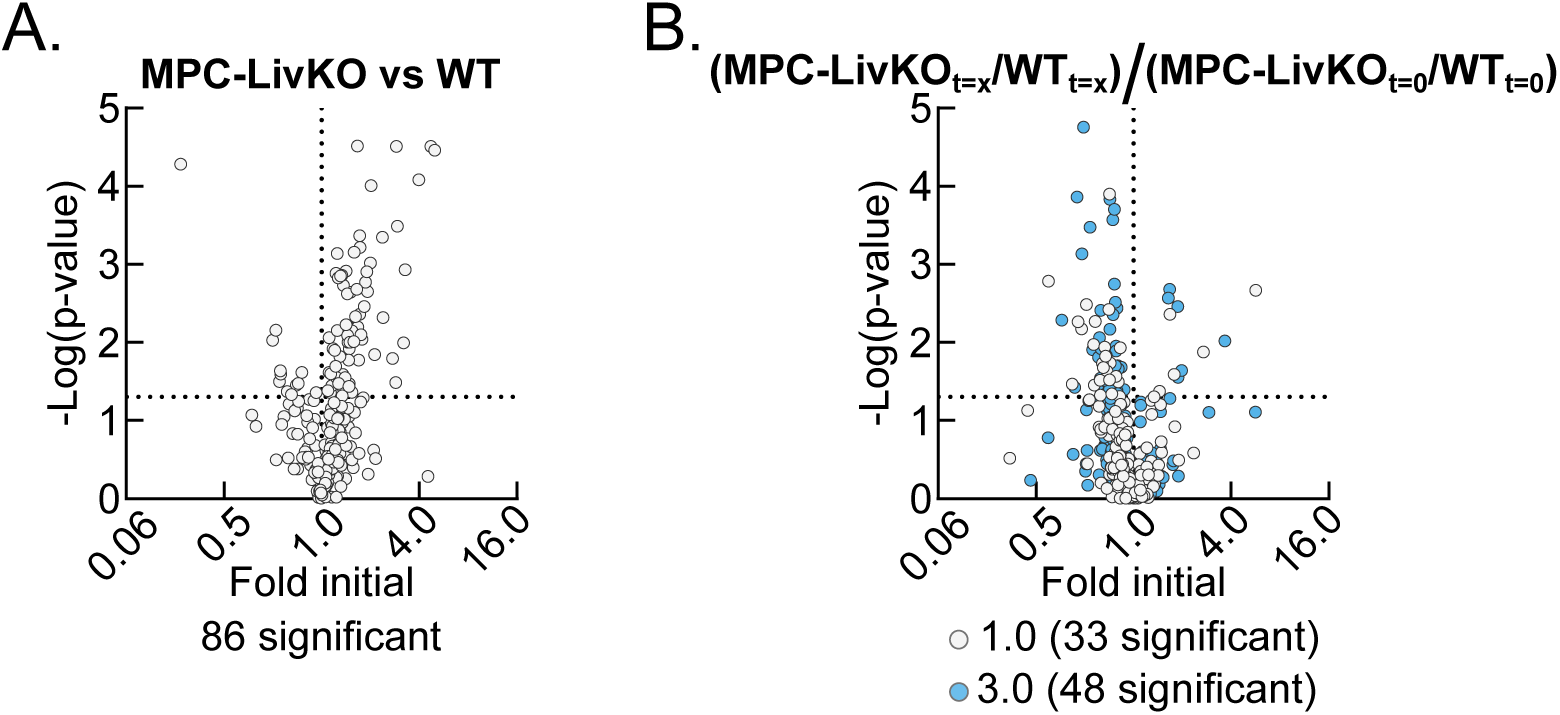
Post-dissection hypoxia results in loss of phenotypic differences. (A) Volcano plot of t0 liver MPC-LivKO versus WT metabolomic data. P-value were calculated using the Student’s t-test on log transformed data. (n=4-5 biological replicates) (B) Volcano plot of t=1 (gray) and t=3 (blue) liver MPC-LivKO/WT versus t=0 MPC-LivKO/WT. Transformed p-values were calculated by one way ANOVA with post-hoc Holm-Sidak multiple comparison test versus t=0 on log transformed data. (n=4-5 biological replicates)

